# Structural basis of the excitatory amino acid transporter 3 substrate recognition

**DOI:** 10.1101/2024.09.05.611541

**Authors:** Biao Qiu, Olga Boudker

## Abstract

Excitatory amino acid transporters (EAATs) reside on cell surfaces and uptake substrates, including L-glutamate, L-aspartate, and D-aspartate, using ion gradients. Among five EAATs, EAAT3 is the only isoform that can efficiently transport L-cysteine, a substrate for glutathione synthesis. Recent work suggests that EAAT3 also transports the oncometabolite R-2-hydroxyglutarate (R-2HG). Here, we examined the structural basis of substrate promiscuity by determining the cryo-EM structures of EAAT3 bound to different substrates. We found that L-cysteine binds to EAAT3 in thiolate form, and EAAT3 recognizes different substrates by fine-tuning local conformations of the coordinating residues. However, using purified human EAAT3, we could not observe R-2HG binding or transport. Imaging of EAAT3 bound to L-cysteine revealed several conformational states, including an outward-facing state with a semi-open gate and a disrupted sodium-binding site. These structures illustrate that the full gate closure, coupled with the binding of the last sodium ion, occurs after substrate binding. Furthermore, we observed that different substrates affect how the transporter distributes between a fully outward-facing conformation and intermediate occluded states on a path to the inward-facing conformation, suggesting that translocation rates are substrate-dependent.

## Introduction

EAATs belong to the Solute Carrier 1 (SLC1) family uptake substrates into cells against their concentration gradients by symporting them with three sodium ions (Na^+^) and a proton (H^+^) and counter-transporting a potassium ion (K^+^)^1–3^. There are 5 EAAT subtypes in humans, sharing similar molecular mechanisms but expressed in different tissues and cell types^4^. EAAT1 and EAAT2 are the principal glial glutamate transporters, with EAAT2 responsible for the uptake of up to 80-90% of the neurotransmitter into astrocytes following rounds of synaptic transmission^5^. EAAT4 and EAAT5 are expressed in Purkinje cells of the cerebellum and retina; they display lower glutamate transport but higher chloride conductance ability^6,7^. By contrast, EAAT3 is expressed in neurons throughout the brain and peripheral tissues, such as epithelial cells of the intestine and kidney and endothelial cells of capillaries^8^. All EAATs can uptake L-Glu, L-Asp, and D-Asp. L-Glu is the brain’s most abundant free amino acid; it mediates transmission at most fast excitatory synapses and is a metabolic hub linking energy metabolism and amino acid biosynthesis in neurons^9^. Under normal conditions, most L-Glu is sequestered inside brain cells, and its excess in the extracellular space can lead to excitotoxicity. L-Asp also fits the criteria of an excitatory neurotransmitter because it excites the NMDA subtype of ionotropic glutamate receptors^10^, but its role in neurotransmission has been questioned^11^. D-Asp, found in the brain and neuroendocrine tissues, shows neuromodulatory activity and may also be a neurotransmitter^12,13^. It is present in high concentrations in the mammalian brain during development but drops sharply postnatally.

EAAT3 is the only EAAT subtype able to transport L-Cys efficiently^14,15^. Neutral SLC1 amino acid transporters (Alanine, Serine, Cysteine Transporters, or ASCTs) can also transport L-Cys^16,17^, while system xc-transporter from the SLC7 family exchanges oxidized L-cystine for glutamate^18^. These transporters are enriched in astrocytes^19–21^, whereas EAAT3 mediates about 90% of L-Cys uptake into neurons^22,23^. In so doing, EAAT3 protects them from oxidative stress because L-Cys is a rate-limiting precursor for antioxidant glutathione (GSH) synthesis. Cysteine is also a substrate for producing the gaseous signaling molecule hydrogen sulfide (H_2_S), a substrate for the post-translational persulfidation of cysteine residues. This evolutionarily conserved modification protects proteins from oxidative stress and can extend the organism’s life^24,25^. EAAT3 deficiency may contribute to a plethora of neurologic pathologies, including ischemic stroke, epilepsy, Parkinson’s, Huntington’s, and Alzheimer’s diseases^26^. Indeed, decreased levels of GSH, present in 2-3 mM concentration in the healthy brain, are an early biomarker of brain aging and Parkinson’s disease^27^. Furthermore, inhibition of EAAT3 by morphine decreases the cell methylation potential and DNA methylation, leading to epigenetic changes implicated in morphine addiction^28^.

EAAT3-mediated L-Glu and L-Asp uptake outside the central nervous system promotes metabolic activity, and the amino acids serve as nucleotide precursors^29^. EAAT3 is also required for rapid metabolic reprogramming in activated B cells^30^ and cancer cells^31^. Recently, EAAT3 has been identified as the "oncometabolite" R-2-hydroxyglutarate (R-2HG) transporter^32^. Tumor cells produce and secrete R-2HG, which acts as a signaling molecule on the surrounding cells, modulating the tumor microenvironment^33^ and might enter endothelial cells via EAAT3, stimulating angiogenesis.

EAAT3 is a homotrimer, with each protomer comprised of the central trimeric scaffold and peripheral transport domains. During uptake, the transport domain undergoes ∼15 Å transmembrane movement combined with a rotation alternating between the outward- and inward-facing states (OFS and IFS); the scaffold domain remains mostly immobile^34,35^. All SLC1 family proteins^36–45^ and its archaeal homologues^46–52^ share this elevator mechanism. A substrate molecule, three Na^+^ ions, and a proton bind to the transport domain in the OFS and dissociate in the IFS; a K^+^ ion binds instead to the IFS and dissociates from the OFS to complete the cycle. The first cryo-EM study on the glycosylation mutant of human EAAT3, hEAAT3g, revealed that the transporter preferentially resided in the IFS in the presence of saturating Na^+^ concentrations^35^. L-Asp showed a very low affinity for the IFS and a greater affinity for the OFS; therefore, we observed growing populations of L-Asp-bound OFS in increasing L-Asp concentrations. In contrast, IFS remained substrate-free. To increase the population of the OFS and observe a lower affinity L-Glu binding, we developed a crosslinking protocol constraining a double cysteine K269C/W441C mutant of EAAT3g in the OFS (hEAAT3-X). The crosslinked protein showed a mixture of the OFS and an atypical intermediate outward-facing state (iOFS*), in which the transport domain moves closer to IFS. The intermediate state exhibited a higher substrate affinity, with L-Glu favoring iOFS* over OFS^34^.

Here, we used hEAAT3-X to examine the structural basis of how EAAT3 recognizes diverse substrates. We combined these studies with ligand-mediated thermal stabilization experiments on hEAAT3g to probe substrate binding in solution and solid-supported membrane (SSM) electrophysiology to test substrate transport. The substrates showed thermal stabilization of the transporters in the order L-Asp > D-Asp > L-Glu >L-Cys, which likely reflects how tightly they bind. Notably, L-Cys showed thermal stabilization only at elevated pH, suggesting it binds in the thiolate form. We observed no hEAAT3 stabilization by R-2HG. SSM electrophysiology showed transport currents for Asp, Glu, and L-Cys, while R-2HG produced no currents. CryoEM imaging of hEAAT3-X in the presence of L-Asp and D-Asp showed transporters predominantly in iOFS* and bound to the amino acids. In contrast, hEAAT3-X, in the presence of R-2HG, pictured the transporter in OFS with an empty and open substrate-binding site, consistent with the biophysical results suggesting that R-2HG is not a transported substrate. Imaging hEAAT3-X in the presence of L-Cys revealed an ensemble of OFS, iOFS*, and a slightly shifted iOFS. The iOFS and iOFS* featured the full complement of bound L-Cys and symported ions. In contrast, OFS, while bound to L-Cys and two Na^+^ ions (at Na1 and Na3 sites), featured a semi-open extracellular gate (helical hairpin 2, HP2) and a disrupted Na2 site. Our work provides the structural basis of promiscuous substrate recognition by EAAT3 and suggests that the substrate binding occurs before the last Na^+^ bounding at the Na2 site and the coupled gate closure.

## Results

### Purified hEAAT3g binds and transports diverse substrates

To compare the binding of different substrates to hEAAT3, we purified the transporter and measured its temperature-induced denaturation in the absence and presence of substrates (**Fig. 1a-c**). hEAAT3g in 200 mM NaCl at pH 7.4 denatured at 69.2 ± 0.2 °C. Additions of 10 mM L-Asp, D-Asp, and L-Glu increased the denaturation temperature by 3.8±0.1, 2.4±0.2, and 1.0±0.1 °C, respectively. In contrast, 10 mM L-Cys, 10 mM D-Glu, or R-2HG did not significantly stabilize the transporter, suggesting that they bind weaker or not at all (**Fig. 1b, c**). To test L-Cys and R-2HG further, we increased their concentrations to 100 mM at pH 7.4 and 8.8 for L-Cys. We observed no significant stabilization by either substrate at pH 7.4. However, at pH 8.8, L-Cys stabilized the transporter by 4.2 ± 0.6°C (**Fig. 1c**). These data suggest that L-Cys binds to the transporter as thiolate. Surprised by the apparent lack of R-2HG binding, we tested whether hEAAT3g reconstituted into liposomes transported R-2HG in solid-supported membrane electrophysiology (SSME). R-2HG carries one less positive charge than L-Glu and D-Glu, but its transport should result in a net uptake of one positive charge and be electrogenic.

**Figure 1.**
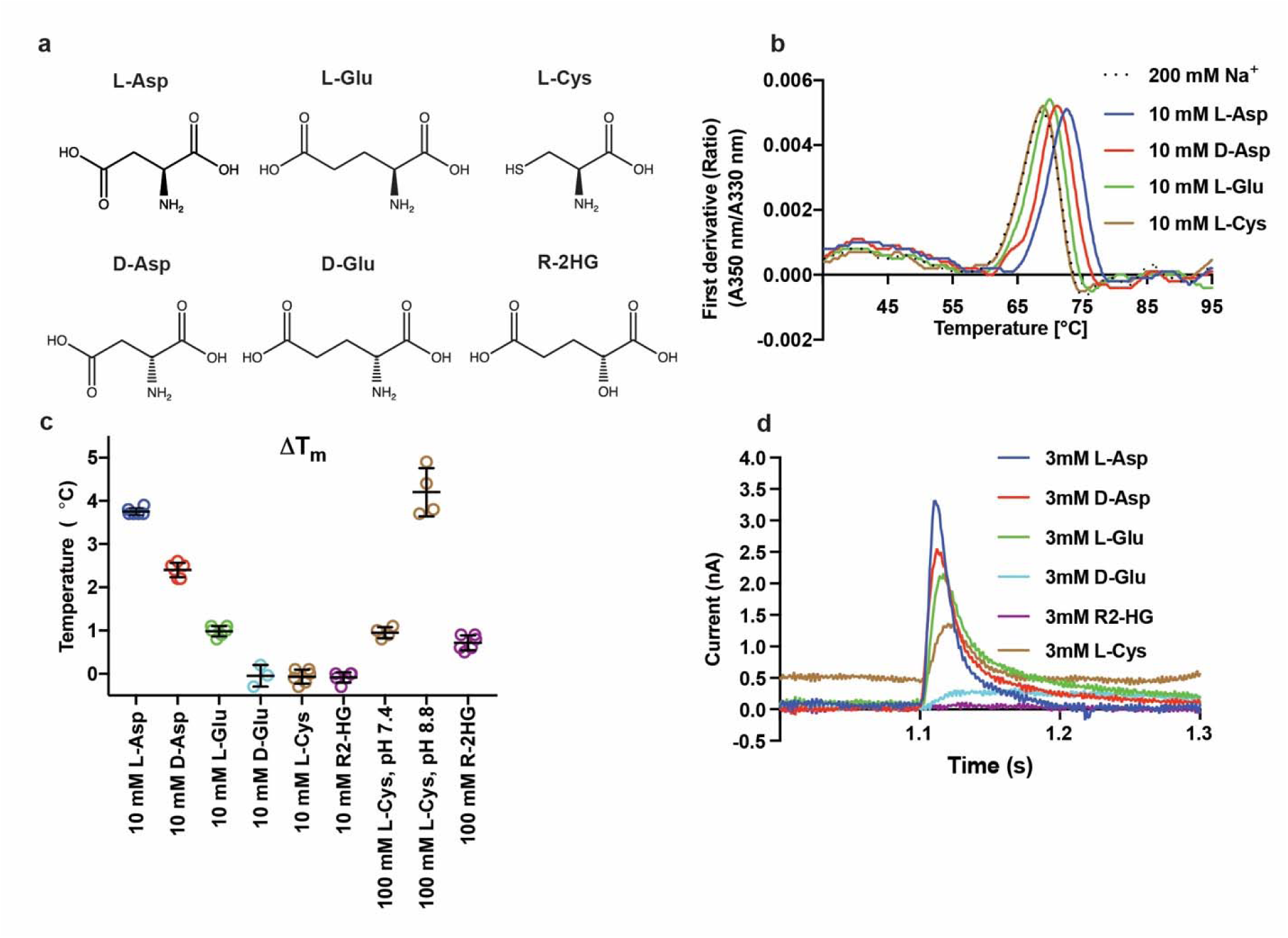
Ligand-dependent thermal stability and transport activity of hEAAT3g. (**a**), Chemical structures of EAAT3 amino acid substrates and R-2HG. (**b**), Representative melting curves of hEAAT3g in 200 mM NaCl (dotted line) and the presence of amino acids, as indicated next to the graph. Shown are the first derivatives of the fluorescence emission intensity ratio at 350 and 330 nm (A_350_/A_330_), with peaks corresponding to the inflections of the sigmoidal melting curves and termed melting temperatures (Tm) (**c**), Tm increases (ΔTm) in the presence of potential substrates compared to NaCl alone. The results for two independent protein preparations (except for D-Glu, which was prepared once), each with multiple technical repeats, are shown; the error bars are the standard deviations. (**d**), Examples of SSME-measured transient currents when immobilized hEAAT3g proteoliposomes were perfused with 3 mM of potential substrates. All experiments were performed using two independent protein purification and reconstitutions, and at least three sensors were used to measure each reconstitution. The color scheme is the same in (**b-d**): L-Asp, blue; D-Asp, red; L-Glu, green; D-Glu, cyan; L-Cys, brown; R-2HG, purple.

Nevertheless, we observed no capacitance peaks upon perfusion of R-2HG. In contrast, perfusion of L- and D-Asp, L-Glu, and L-Cys over the same SSM chip produced robust peaks, and perfusion of D-Glu produced a small but reproducible capacitance current (**Fig. 1d**).

### Structures of hEAAT3-X bound to substrates

To examine substrate binding structurally, we introduced K269C/W441C into Cysmini EAAT3 as previously described^34,53,54^. Hg^2+^-mediated cross-linking traps the transporter in iOFS*, iOFS, and OFS (hEAAT3-X), which show high-affinity L-Asp and L-Glu binding and are ideal for examining varying potential substrates. Following cross-linking, we purified hEAAT3-X by SEC in 100 mM NMDG-Cl (Apo condition), split the eluted protein into two samples, and supplemented them with 200 mM NaCl and 10 mM L-Asp or R-2HG before freezing cryo-EM grids. Data processing on the L-Asp sample yielded a well-resolved map at 2.87 Å resolution. The map revealed iOFS* conformation with a closed substrate gate (helical hairpin 2, HP2) and a well-resolved density corresponding to the bound L-Asp (**Fig. 2a, c, Supplementary Fig. 1, Supplementary Table 1**); we found no additional minor conformations in 3D classifications. In contrast, the R-2HG dataset yielded a 3.07 Å resolution OFS map featuring a wide-open HP2 gate, nearly identical to the OFS observed in Na^+^ buffers without substrates (**Fig. 2b, d, Supplementary Fig. 2, Supplementary Table 1**). We found 8 % protomers in iOFS with no density corresponding to R-2HG (**Supplementary Fig. 2c)**; this conformation is nearly identical to the minor state observed in Na^+^ buffer without substrate^34^.

**Figure 2.**
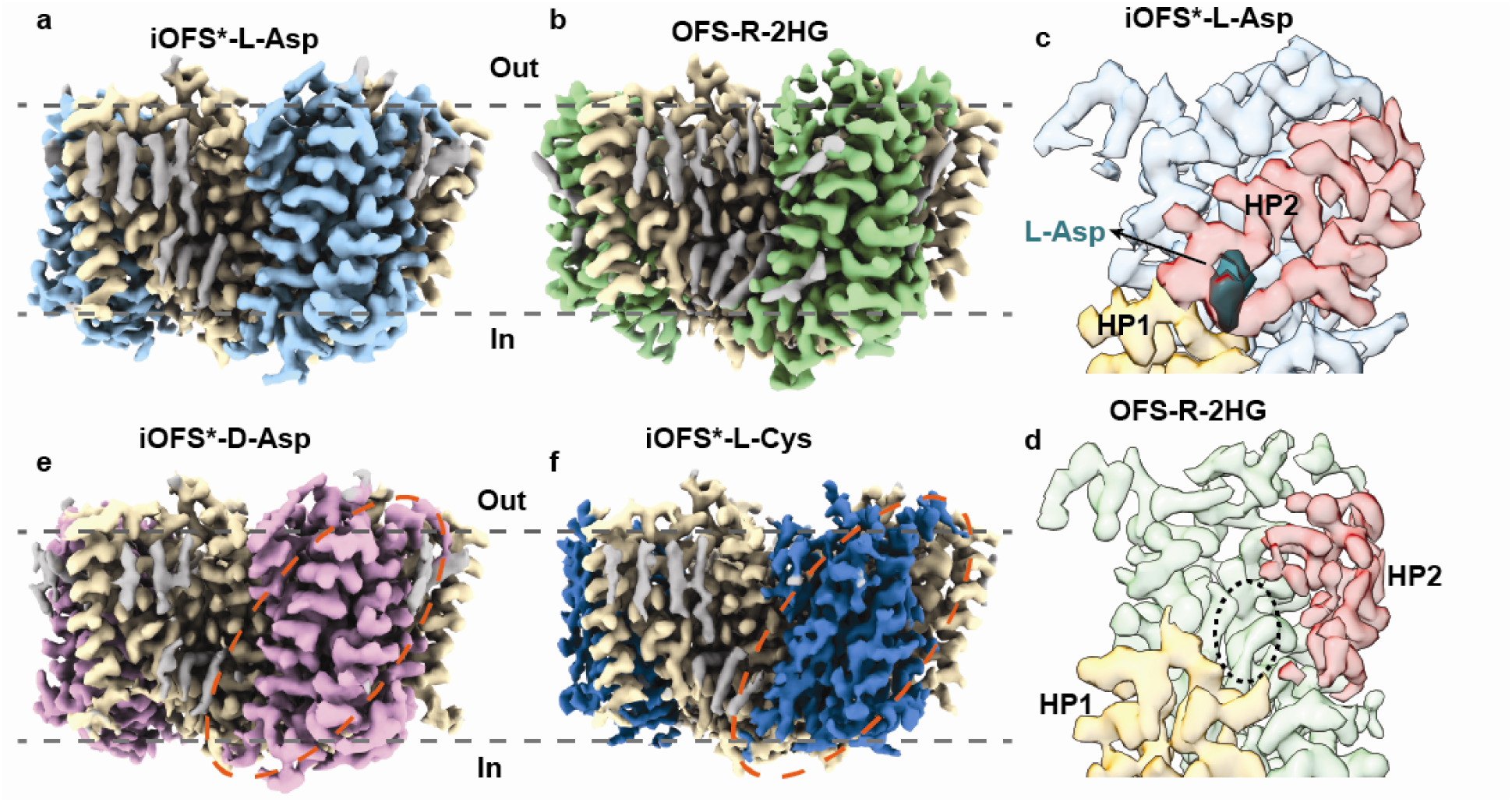
The structures of hEAAT3-X with 10 mM substrates. The overall structure of hEAAT3-X with 10 mM L-Asp (**a**), R-2HG (**b**), D-Asp (**e**), or L-Cys (**f**); The orange dashed ovals highlight the transport domain density of iOFS*-D-Asp and iOFS*-L-Cys. The scaffold domains are colored in wheat, the lipid densities are gray, and the transport domains are multicolored with L-Asp, light blue; R-2HG, green; D-Asp, pink; and L-Cys, dark blue. (**c, d**), The structures of iOFS*-L-Asp (**c**) and OFS-R2HG (**d**) transport domains. Helical hairpin 1 (HP1) and HP2, which define the location of the substrate-binding site, are colored yellow-orange and red, respectively. HP2 of iOFS*-L-Asp is closed, with the bound L-Asp colored in teal (**c**); The HP2 of OFS-R2HG is wide open, and the ligand-binding cavity, emphasized by the black dotted oval, is empty (**d**). The contour levels of the iOFS*-L-Asp, OFS-R-2HG, iOFS*-D-Asp, and iOFS*-L-Cys trimer maps are 0.614, 0.34, 0.614, and 0.62, respectively; the gray dashed lines represent an approximate position of the lipid bilayer.

Next, we prepared another batch of apo hEAAT3-X, which we supplemented with 200 mM NaCl and 10 mM L-Cys or D-Asp. Because L-Cys can break Hg^2+^-mediated cysteine crosslink, we rapidly mixed ice-cold EAAT3-X with L-Cys and froze grids immediately, in less than 10 seconds. Processing of the D-Asp dataset produced a 2.73 Å resolution density map with resolved scaffold and transport domains corresponding to iOFS* (**Fig. 2e, Supplementary Fig. 3, Supplementary Table. 1**), and 3D classification did not reveal the presence of any other states. Interestingly, we previously found that for hEAAT3-X bound to L-Glu, about 14% of protomers were in the OFS conformation, with the remainder in iOFS*. In contrast, we found no OFS structural classes in the current L-Asp or D-Asp datasets. Thus, we hypothesize that ligands can affect the transport domain distribution of the EAAT3-X.

### Conformational ensemble of L-Cys-bound hEAAT3-X

Processing the L-Cys dataset yielded a density map at 2.36 Å resolution with applied C3 symmetry. The map showed a well-resolved scaffold domain density but a blurred transport domain density (**Fig. 2f, Supplementary Fig. 4**). Because we observed no such blurring in the D-Asp dataset, which was prepared simultaneously, we reasoned that it was not due to damaged protein and might reflect protein dynamics. To uncover the complete conformational ensemble of L-Cys-bound EAAT3-X, we performed symmetry expansion and optimized the parameters of the local 3D classification in Relion^55^. When the class number, *K*, and the regularization parameter, *T*, were set to 20 and 40, we identified 4 distinct structural classes. Further local refinement produced EM maps corresponding to OFS, iOFS, iOFS*, and IFS with resolutions of 2.58, 2.99, 2.60, and 2.94 Å. (**Fig. 3, Supplementary Figs. 5, 6, Supplementary Table 1**). The IFS presence indicates that the Hg^2+^ crosslink is disrupted in a fraction of EAAT3-X molecules during grid preparation. Aided by the substantial number of expanded particles (3.3 million), the EM map of the lowly populated iOFS class, comprising 1.8% of particles, is well-resolved. We could not sort out iOFS with smaller *K* values, such as 5 and 10.

**Figure 3.**
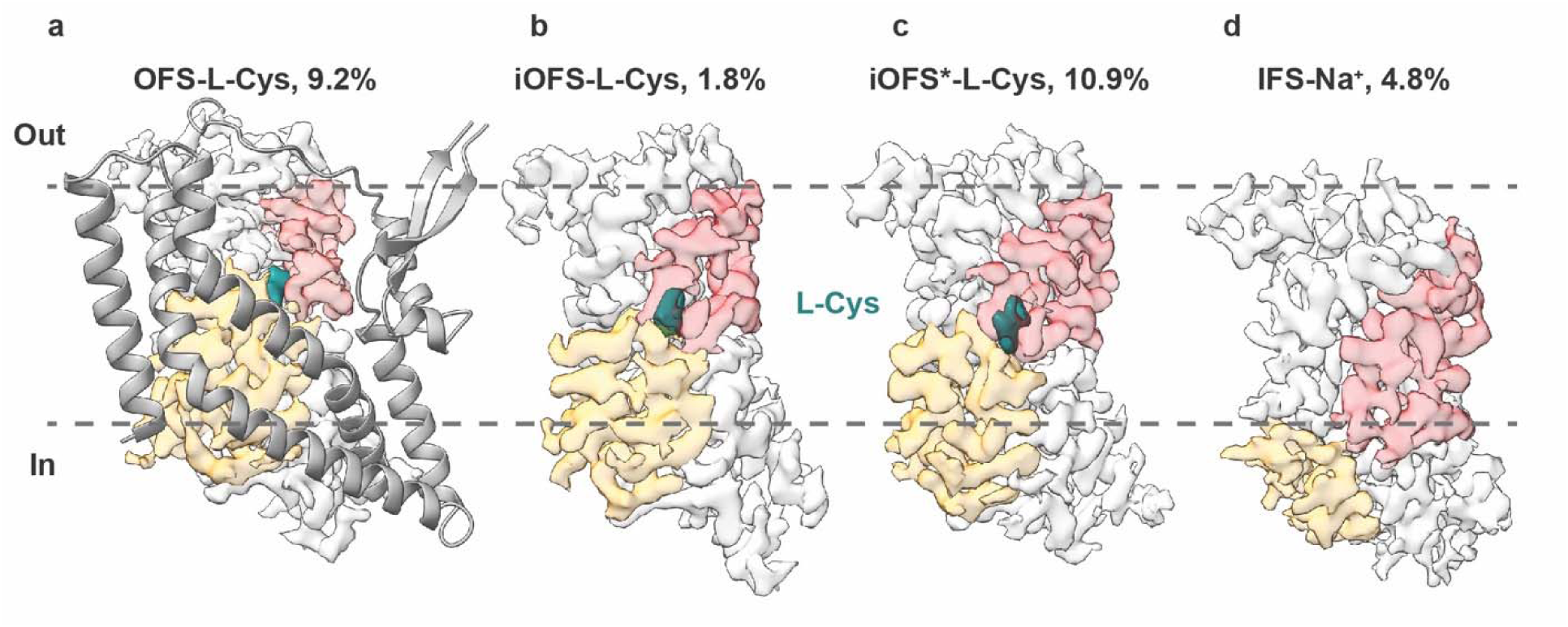
Conformational ensemble of hEAAT3-X with 10 mM L-Cys. (**a**), The overall structure of OFS-L-Cys. The scaffold domain is colored in gray and shown as a cartoon; the transport domain is colored in light gray, with the HP1 and HP2 colored in yellow-orange and red, respectively; the density of L-Cys is colored in teal. The transport domains of iOFS-L-Cys (**b**), iOFS*-L-Cys (**c**), and IFS-Na^+^ (**d**) are colored as in (**a**). For clarity, their scaffold domains, which were aligned to OFS-L-Cys, are not shown. The contour levels of these maps are 0.65, 0.54, 0.61, and 0.43, respectively.

We observed strong non-protein density in the substrate-binding pocket of OFS, iOFS, and iOFS* maps, which we modeled as L-Cys (**Fig. 3a-c**). In contrast, there was no ligand density in the IFS map (**Fig. 3d**). Furthermore, the HP2 gate in the IFS map is wide open, suggesting that it is bound to Na^+^ ions only, consistent with the low substrate affinity of the IFS we previously reported for hEAAT3g^35^. The overall structure of L-Cys-bound iOFS* (iOFS*-L-Cys) is remarkably similar to iOFS*-L-Glu; the RMSD calculated by the whole structure alignment is 0.628 Å. The superposition of iOFS-L-Cys and iOFS*-L-Cys aligned on the scaffold domain shows that the iOFS-L-Cys transport domain is positioned more outward than in iOFS*. It corresponds more closely to the iOFS observed in potassium-bound EAAT3-X, iOFS-K^+34^ (**Supplementary Fig. 7**).

### Structural basis of ligands recognition by EAAT3

The iOFS*-Cys structure shows that L-Cys is coordinated identically to L-Glu. Its main chain carboxylate interacts with the sidechain of N451 in TM8 and the main chain and sidechain oxygens of S333 in HP1, and its amino group interacts with the sidechain of D444 in TM8. The L-Cys sidechain sulfur atom is 2.9 Å away from the guanidinium group of R447 (**Fig. 4b, d**), which typically coordinates the sidechain carboxylate of L-Glu, consistent with the bound L-Cys being in thiolate form. Further comparison between EAAT3-X bound to L- and D-Asp, L-Glu, and L-Cys shows that the R447 sidechain moves slightly outward and assumes a different rotamer in the L-Glu- and L-Cys-bound structures compared with the L-Asp- and D-Asp-bound conformations (**Fig. 4**). The superposition of EAAT3-X substrate-binding pockets shows that L-Glu, L-Cys, and L-Asp bind to EAAT3 in similar poses with their amino groups pointing toward HP2 and interacting with D444. In contrast, the D-Asp’s amino group points toward TM8 while still interacting with D444 (**Fig. 4**). The subtle binding pose difference between L- and D-Asp is consistent with the previous structural study on Glt_Tk_^56^. Thus, EAAT3 recognizes diverse substrates by fine-turning sidechain conformations in the binding pocket and subtle changes in the substrate poses.

**Figure 4.**
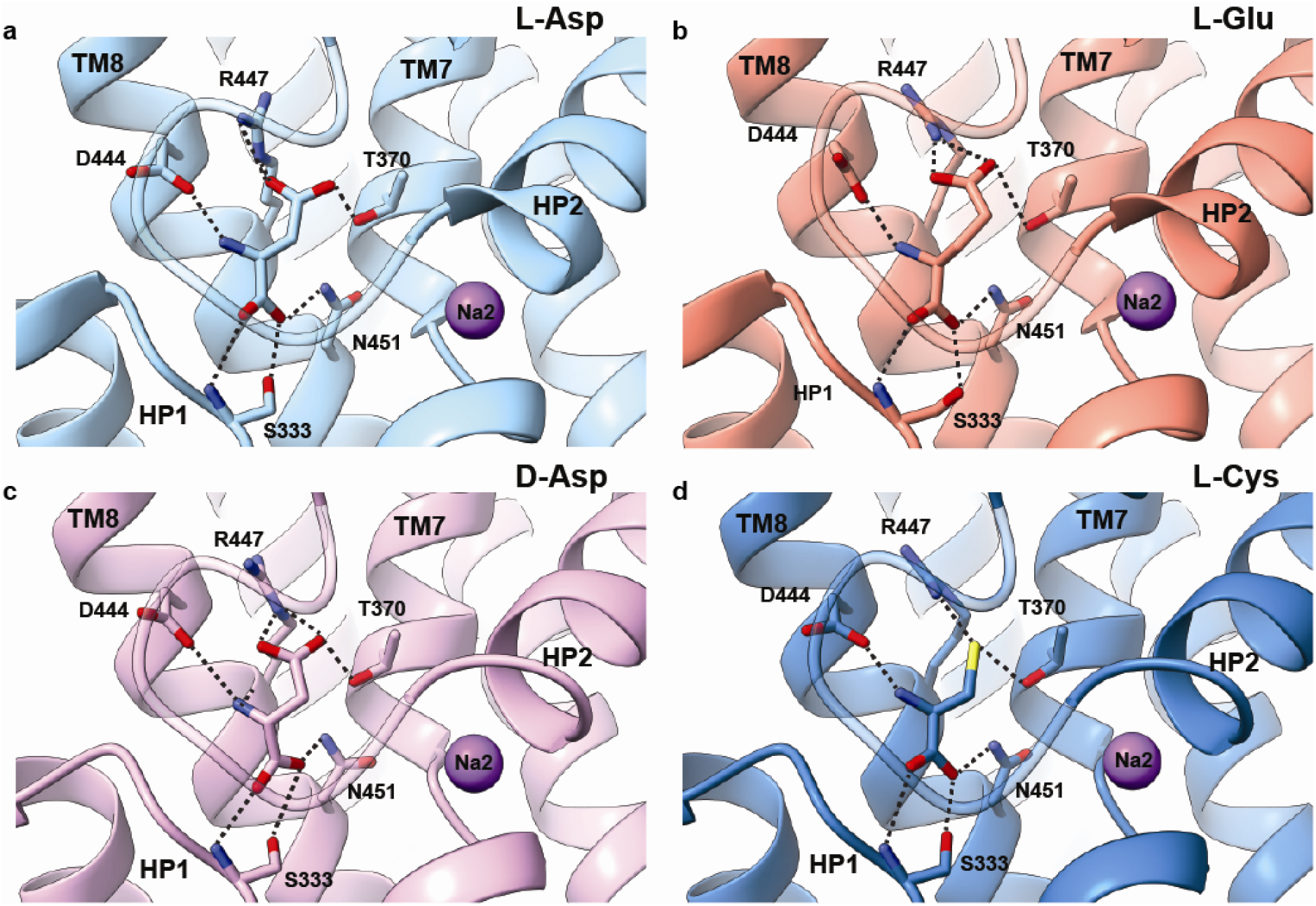
The substrate-binding pocket of hEAAT3-X with different substrates. Binding pockets with L-Asp (**a**), L-Glu (**b**, PDB: 8CTC), D-Asp (**c**), and L-Cys (**d**). The substrates and interacting residues are shown as sticks. Dashed black lines show the interactions between the residues and the substrates. The transport domains are superposed on their cytoplasmic halves (residues 314-372 and 442-465).

### Partially open gate in the outward-facing L-Cys-bound state

HP2 gate occludes substrates in the binding site of EAATs before their translocation across the membrane. The superposition of the transport domains (residues 80-120 and 280-470) of L-Cys-bound iOFS* and iOFS with OFS produced RMSDs of 0.607 Å and 0.692 Å, suggesting that overall transport domains are almost identical in the three states. We found well-defined density at the three sodium sites in iOFS*, and the surrounding residues feature appropriate geometry to coordinate Na^+^ (**Supplementary Fig. 8a**). Thus, iOFS*-L-Cys is in the fully-bound occluded state with L-Cys, three Na^+^ ions, and a closed HP2. iOFS shows nearly identical geometry of the sodium-binding sites, an excess density corresponding to L-Cys, and a closed HP2, suggesting it is also a fully-bound occluded state, even if the resolution is insufficient to visualize Na^+^ ions unambiguously. By contrast, in OFS, we could find extra densities at the substrate-binding site, the Na1 and Na3 sites, but not the Na2 site. The HP2 tip (i.e., the GVPN_410-413_ loop between the two helical arms of HP2) is positioned roughly in the middle between the wide-open OFS-Na^+^ state and the fully-bound, closed iOFS*-L-Cys state; it moves away from the substrate-binding pocket by about 4.5 Å compared to the iOFS* structure (**Fig. 5a**). Thus, the substrate-binding pocket is exposed to solvent (**Fig. 5b, c**), and we found two extra densities assigned to water molecules in the pocket. While OFS-L-Cys lacks interactions between L-Cys and HP2, which are present in iOFS*-L-Cys (**Supplementary Fig. 8c, d**), the remainder of L-Cys coordination is preserved (**Supplementary Fig. 8e, f**). In the iOFS*-L-Cys structure, residues SASIGA_403-408_ form the last 2 helical turns of the HP2a arm, and the main chain oxygen atoms of S405, I406, and A408 coordinate the Na^+^ at the Na2 site with the sulfur of M367 and main chain oxygen of T364 in TM7a (**Fig. 5d, e**). The sidechain of the conserved S405 residue points toward TM7a, forming a water-mediated hydrogen bond and stabilizing the closed HP2 configuration. In contrast, the SASIGA_403-408_ region is unwound in the OFS-L-Cys structure; the S405 side chain weakly interacts with the L-Cys thiolate group (**Supplementary Fig. 8c, d**). The geometry of the Na2 site is disrupted with distances to the main chain oxygen atoms of S405, I406, and A408 being 1.3, 3.8, and 5.5 Å, respectively (**Fig. 5f, supplementary Fig. 8a, b**). These features suggest the OFS-L-Cys structure captures an intermediate before the last sodium binds at the Na2 site and the HP2 gate closes (**Supplementary Movie 1**).

**Figure 5.**
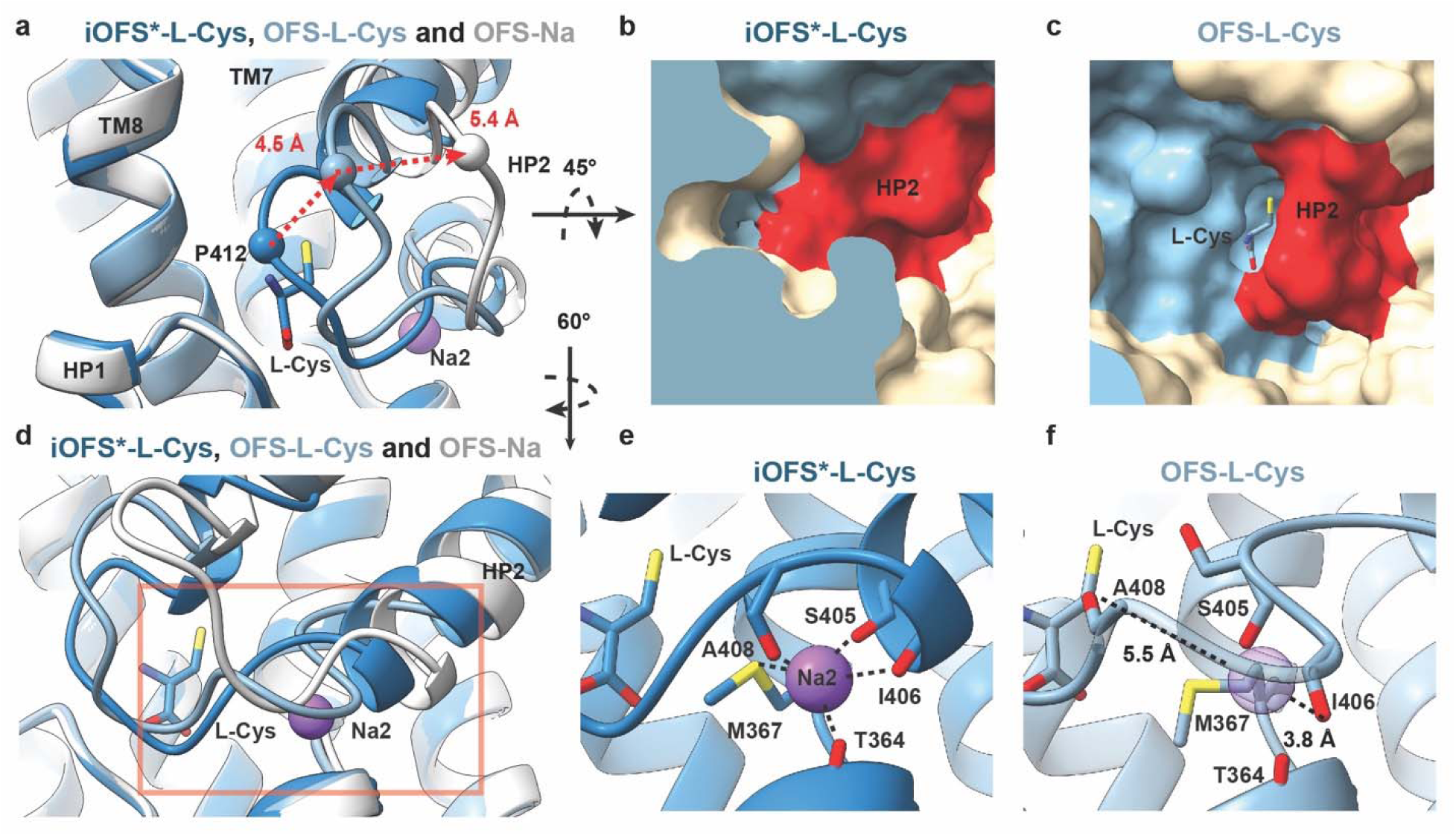
The partially open HP2 gate in OFS-L-Cys. (**a**), The tip of HP2 of OFS-L-Cys (pastel blue) is in between the wide-open HP2 observed in Na^+^-only bound OFS (white, PDB: 8CV2) and the fully closed HP2 in iOFS*-L-Cys (slate blue). The distances between α carbons of P412 in the HP2 tip of the three states are shown as dashed red lines. The transport domains are superposed as in Figure 4. Only L-Cys in iOFS* is shown as sticks for clarity. (**b, c**), The surface representation of iOFS*-L-Cys (**b**) and OFS-L-Cys (**c**) binding sites. HP2 (red) occludes the pocket in iOFS* (**b**) but allows solvent access in OFS (**c**). (**d**), The Na2 site in the three states with protein structures colored as in (**a**). The red box shows the part of the structure enlarged in **e** and **f**. (**e, f**), The formed Na2 site with the bound Na^+^ ion in iOFS*-L-Cys (**e**), and the distorted Na2 site in OFS-L-Cys (**f**). The dashed black lines represent the interactions between residues and the ion (**e**) or the distance between the main chain oxygens of I406 and A408 and the site of Na2 binding, shown as a transparent purple sphere (**f**).

## Discussion

EAAT3, an electrogenic acidic amino acid and cysteine transporter, orchestrates amino acid metabolism and protects cells from oxidative stress. Our structures visualize hEAAT3 recognizing four substrates: L-Asp, D-Asp, L-Glu, and L-Cys. Supported by the binding assays, they suggest that EAAT3 transports L-Cys in thiolate form, consistent with previous studies^14^. The transporter coordinates acidic amino acids and L-Cys thiolate by fine-tuning the position of the same residues, especially the pivotal R447, which coordinates the substrate side-chain acidic moiety. R447 is replaced with threonine and cysteine in the neutral amino acid transporters ASCT1 and 2, respectively, and a recently reported structure of ASCT2 with L-alanine^45^ suggests that ASCT2 transports L-Cys in the thiol form. (**Supplementary Fig. 9a, b**). The EAAT3 R447C mutant does not bind or transport acidic amino acids while it still transports L-Cys and neutral amino acids via the electroneutral exchange mechanism, similar to ASCT2^57^. The main chain amino and carboxyl groups or the substrate are coordinated by the highly conserved D444 and N451, respectively. Together, D444, R447, and N451 are the critical determinants of substrate specificity.

R-2HG is an oncometabolite that rewires the metabolism of cancer cells by inhibiting α-KG-dependent dioxygenases and changing epigenetic modification patterns^58^. R-2HG might also promote tumor growth through other mechanisms^59,60^. Recently, it was proposed that R-2HG enters cells and their mitochondria through EAAT3 localized to the plasma and mitochondrial membranes, respectively^32^. This proposal prompted us to examine R-2HG binding and transport using purified protein. We found that up to 100 mM R-2HG did not significantly thermally stabilize hEAAT3g in differential scanning fluorimetry experiments, suggesting that it binds weakly or does not bind. The SSME assays performed with 3 mM substrates, a saturating concentration for L-Asp, showed similar transport currents for L-Asp, D-Asp, and L-Glu and a smaller current for L-Cys (**Fig. 1c**). The D-Glu transport current was shallow, persisting much longer ligand perfusion time, suggesting that D-Glu transport is very slow. Indeed, D-Glu is a low-affinity EAAT3 substrate with Km of ∼1.8 mM, approximately 60-fold higher than L-Glu^61^. In contrast, R-2HG produced no current (**Fig. 1c)**. Finally, R-2HG added at 10 mM did not bind to EAAT3-X in cryo-EM imaging experiments. R-2HG is an analog of D-Glu, in which an alcohol moiety replaces the amino group. Compared to D-Glu, R-2HG loses a critical salt bridge between the amino group and D444. Mutations of D444 in EAAT3 cause a dramatic reduction of affinity for amino acids^62,63^, suggesting that EAAT3 would bind R-2HG even weaker than D-Glu. Thus, our results and structural considerations do not support the hypothesis that EAAT3 is the R-2HG transporter in cancer cells. However, it should be noted that R-2HG concentrations in tumors can reach 30 mM^60^, and it is, in principle, possible that EAAT3 transports R-2HG with very low affinity.

L-Cys is a rate-limiting substrate of GSH biosynthesis and, therefore, is an important metabolite in maintaining the cell redox status, methylation potential, and protection against oxidative stress in all cell types. In the bloodstream, ∼95 % of L-Cys is oxidized to cystine, which can be taken up by SLC7A11 transporter system xc-into glial cells and reduced to L-Cys. Interestingly, ASCT2, which could also contribute to L-Cys uptake into glia, has a similar Km of ∼20 μM for L-Cys and other neutral amino acids but a nearly 10-fold lower Vmax, suggesting L-Cys is not an efficient substrate^17^. EAAT2, highly expressed in glial cells, does not uptake L-Cys well because of its low affinity for the amino acid with Km of ∼1-2 mM, much higher than ∼250 µM concentration of L-Cys and its derivatives in the plasma^64^. EAAT3 is the main L-Cys transporter in the neurons with a Km of ∼100-200 µM^23^, about 10-fold above L-Glu, and a similar Vmax. Interestingly, the comparison between L-Glu-bound EAAT3 and EAAT2 and L-Cys-bound EAAT3 does not reveal significant structural differences between EAAT2 and EAAT3 that would explain similar affinity for L-Glu and drastically different affinities for L-Cys. Thus, allosteric effects outside of the binding site might contribute to different substrate specificities. Indeed, previous studies in an archaeal homolog Glt_Ph_ suggested that differences in protein packing and dynamics might contribute to substrate affinity and selectivity ^65,66^.

Kinetic studies on EAATs and their homologs suggest that substrate and ion binding proceeds via partially bound intermediates, such as the transporter bound to the substrate and one or two sodium ions, before forming the transport-component complex of the substrate and three sodium ions. EAATs bind substrates rapidly on the sub-millisecond time scale but transport them slower, with turnover times estimates in milliseconds to tens of milliseconds, resulting in biphasic electrical currents comprised of the binding peak currents and the lower steady-state currents^67,68^.

The initial binding is weak, with Kd of ∼140 µM for EAAT2 significantly higher than the transporter Km of 10-20 µM^69^. Our structure of EAAT3 in OFS with bound L-Cys and partially open HP2 gate with clear densities at the Na1 and Na3 sites but a distorted empty Na2 site might directly visualize the proposed low-affinity binding intermediate.

Interestingly, the transporter has demonstrated different conformational preferences depending on the substrate. Thus, in L- and D-Asp, we only observed the EAAT3-X in the iOFS*. In contrast, the transporter bound to L-Glu populated iOFS* and OFS with closed HP2, while the transporter bound to L-Cys populated iOFS*, iOFS, and OFS with the partially open HP2 gate. These observations should be taken cautiously because the grids were not prepared identically in all cases: the L-Cys grids were prepared by rapidly freezing the protein seconds after adding the substrate, while others were prepared using protein equilibrated with substrates. Nevertheless, the observed differences suggest that to the relative energies of transporter states along the transport cycle depend on the substrates. If so, we would speculate that the transporters might show substrate-dependent transport rates, as was shown for EmrE^70^.

## Methods

### Protein expression and purification

The hEAAT3g and Cysmini K269C/W441C EAAT3g proteins were purified as previously described. In brief, hEAAT3 constructs were expressed in suspension FreeStyle^TM^ 293-F cells. Isolated membrane pellets were solubilized in a buffer containing 50□mM Tris-Cl at pH□8.0, 1□mM L-Asp, 1□mM EDTA, 1□mM tris(2-carboxyethyl) phosphine (TCEP), 10% glycerol, 1:200 dilution of protease inhibitor cocktail (catalog no. P8340, Sigma-Aldrich), 1□mM phenylmethylsulfonyl fluoride (PMSF), 1% dodecyl-β-D-maltopyranoside (DDM, Anatrace) and 0.2% cholesteryl hemisuccinate (CHS; Sigma-Aldrich) at 4□°C, overnight. The insoluble material was removed by centrifugation, and the supernatant was incubated with Strep-Tactin Sepharose resin (GE Healthcare) for 1□h at 4□°C. The resin was washed with a buffer containing 50□mM Tris-HCl at pH□8.0, 200□mM NaCl, 0.06% glyco-diosgenin (GDN, Anatrace), 1□mM TCEP, 5% glycerol and 1□mM L-Asp (wash buffer). The protein was eluted with a wash buffer supplemented with 2.5□mM D-desthiobiotin (elution buffer). The N-terminal Strep II and GFP tag was cleaved by overnight PreScission protease digestion at 4 °C. hEAAT3g and Cysmini K269C/W441C EAAT3g were purified by size-exclusion chromatography (SEC) in a buffer containing 20□mM HEPES-Tris at pH□7.4, 1□mM L-Asp, and 0.01% GDN with/without 1mM TCEP. The Cysmini K269C/W441C EAAT3g protein was concentrated to ∼0.5 mg/ml and incubated with a 20-fold molar excess of HgCl_2_ for 15 min at room temperature. Then, crosslinked hEAAT3-X was purified by SEC in a buffer containing 20 mM Hepes-Tris pH 7.4, 100 mM N-methyl-D-glucamine (NMDG) chloride, and 0.01% GDN to remove sodium and L-Asp. The eluted protein was diluted ∼1,000-fold into a buffer containing 20 mM Hepes-Tris pH 7.4, 200 mM NaCl, and 0.01% GDN and concentrated to ∼5 mg/ml using 100 kD MWCO concentrators (Amicon). EAAT3-X in 200 mM NaCl was incubated with the final concentration of 10 mM L-Lap, D-Asp, or R2-HG for about 1 hour on ice before making grids. EAAT3-X in 200 mM NaCl was mixed with L-Cys at a final concentration of 10 mM and put on grids immediately.

### Thermostability assays

Purified hEAAT3g was diluted ∼4000-fold in a buffer containing 50 mM Hepes-Tris pH 7.4, 100 mM NMDG, and 0.01% GDN and concentrated to ∼100 µM using a 100 kD MWCO concentrator. The concentrated protein was diluted 20-fold in a buffer containing 50 mM Hepes-Tris pH 7.4, 200 mM NaCl, and 0.01% GDN, supplemented with 10 mM or 100 mM ligands. To promote L-Cys binding, the concentrated protein was diluted 20-fold in a buffer containing 50 mM Tris-Cl, pH 8.8, 200 mM NaCl, and 0.01% GDN, supplemented with 100 mM L-Cys. The thermostability assay was performed using Tycho NT.6 (NanoTemper Technologies). Protein samples were heated from 35 °C to 95 °C at 30 °C per minute; the intrinsic protein fluorescence was recorded at 330 nm and 350 nm. The amplitude ratio, A350/A330 as a function of temperature, and its first derivative were calculated by the Tycho NT.6 software. The inflection temperature (Ti) corresponds to the peak of the derivative. All measurements were repeated at least thrice on independently prepared protein samples except the D-Glu sample.

### Proteoliposome reconstitution and solid-supported membrane electrophysiology (SSME)

The proteoliposome reconstitution and SSME were performed as previously described^34^. In brief, 4 mg/ml liposomes comprising 5:5:2 (w:w) 1-palmitoyl-2-oleoyl-sn-glycero-3-phosphocholine (POPC, Avanti Polar Lipids), 1-palmitoyl-2-oleoyl-sn-glycero-3-phosphoethanolamine (POPE, Avanti Polar Lipids) and CHS were extruded 11 times through 400 nm polycarbonate membranes (Avanti Polar Lipids) in a buffer containing 50 mM Hepes-Tris, pH 7.4, 200 mM NaCl, 1mM TCEP, 1 mM L-Asp. The resulting unilamellar liposomes were destabilized by incubating with 5:1 (w:w) DDM-CHS at a 1:0.75 lipid-detergent ratio for 30 min at 23 °C. 0.4 mg purified hEAAT3g was incubated with liposomes at a lipid-protein ratio (LPR) of 10 for 30 min at 23 °C. The detergent was removed by incubating with 100 mg fresh Bio-Beads SM-2 (Bio-Rad) for 1h at 23 °C, 1 h at 4 °C (three times), overnight at 4 °C, and finally 2 h at 4 °C. The proteoliposomes were collected by centrifugation at 86,600 g for 45 min at 4 °C and were resuspended in the SSME resting buffer containing 100 mM potassium phosphate, pH 7.4, 2 mM MgSO_4_. The proteoliposomes were frozen in liquid nitrogen and thawed at room temperature. The centrifugation and freeze-thaw steps were repeated three times for buffer exchange. Then, the proteoliposomes were extruded 11 times through a 400 nm polycarbonate membrane and immediately deposited onto the SF-N1 sensor 3mm (Nanion Technologies). The transport-coupled currents were recorded on a SURFE2R N1 instrument (Nanion Technologies). The non-activating buffer containing 100 mM sodium phosphate, pH 7.4, and 2 mM MgSO_4_ flowed through the sensor to build ion gradients across the proteoliposomes. The transport-coupled current was activated by flowing the activation buffer containing 100 mM sodium phosphate, pH 7.4, 2 mM MgSO_4_, and 3 mM ligands. At least three sensors were recorded for each independent proteoliposome preparation.

### Cryo-EM sample preparation and data acquisition

3.5 μl of protein samples at ∼5 mg/ml were applied to glow-discharged Quantifoil R1.2/1.3 holey carbon-coated 300 mesh gold grids. The grids were blotted for 3 s and plunge-frozen into liquid ethane using FEI Mark IV Vitrobot at 4°C and 100% humidity. For the hEAAT3-X with 10 mM L-Asp sample, 13,349 movies were collected at a nominal magnification of 100,000-fold with a calibrated pixel size of 1.16 Å. The nominal defocus value −1.0 ∼ −2.5 µm and total dose 40 e^-^/Å^2^ (dose rate 7.98 e^-^/Å^2^/s) were applied to the data collection. For the hEAAT3-X with 10 mM R-2HG dataset, 11,952 movies were collected at a nominal magnification of 105,000-fold with a calibrated pixel size of 0.8443 Å using the counting model. The nominal defocus value of ∼0.8-2.2 µm and the total dose of 50.54 e^-^/Å^2^ (dose rate 33.69 e^-^/Å^2^/s) were applied to data collection. For the hEAAT3-X with 10 mM D-Asp sample, 4,190 movies were collected at a nominal magnification of 64,000-fold with a calibrated pixel size of 1.076 Å using the counting model. A nominal defocus value −0.5 ∼ −2.0 µm was applied to data collection, with the total dose 52.19 e^-^/Å^2^ (dose rate 26.09e^-^/Å^2^/s) distributed over 40 frames in each movie. For data collection on hEAAT3-X with 10 mM L-Cys sample, subset A (5,765 movies) and subset B (3,757 movies) were collected at a nominal magnification of 105,000-fold with a calibrated pixel size of 0.4125 Å using the super-resolution model. A nominal defocus value −0.8∼-2.4 µm was applied to data collection, with the total dose 58.25 e^-^/Å^2^ (dose rate 29.12 e^-^/Å^2^/s, subset A) and 58.01 (dose rate 29.00 e^-^/Å^2^/s, subset B) e^-^/Å^2^ distributed over 50 frames in each movie. The hEAAT3-X with 10 mM L-Asp data was auto-collected using EPU on Glacios with Falcon4i camera at Weill Cornell Cryo-EM facility; other datasets were auto-collected using Leginon^71^ on Titan Krios with Gantan K3 camera at the Simons Electron Microscopy Center (SEMC) at New York Structural Biology Center (SEMC-NYSBC, R-2HG, and D-Asp datasets), and at New York University Langone’s Cryo-EM laboratory (L-Cys dataset) and. All microscopes were equipped with a 20 -eV energy filter.

### Cryo-EM image processing

For the hEAAT3-X with 10 mM L-Asp dataset, the movies were aligned using MotionCorr2^72^ implemented in Relion 4, and the micrograph CTF parameters were estimated using CtfFfind-4.1^73^. Over 12 million particles were selected by Laplacia-of-Gaussian (LoG)^74^ and extracted with a box size of 120 pixels (2-fold binning) from 12,021 micrographs. The particles were divided into four parts and imported into CryoSPARC v4^75^ for 2D classification. 378,103 particles showing clear secondary features were selected and used for 1 round of *ab initio* reconstruction; the resulting 211,611 particles were subjected to nonuniform refinement^76^ (hereafter NUR) with C1 symmetry to generate a good template, while for generating 5 decoy templates, 448,378 junk particles were selected and subjected to *ab initio* reconstruction for less than 10 iterations. More than 10 million particles after 2D selection that removed obvious non-protein junks (2D cleaning) were further cleaned by heterogeneous refinement with 1 good template and 5 decoy noise volumes (heterogeneous refinement cleaning, HRC). 1,240,537 particles were refined to 4.84 Å by NUR with C1 symmetry. Then, the particles were re-imported into Relion through PyEM^77^ and extracted with a box size of 240 pixels without binning. These particles were imported into CryoSPARC and subjected to HRC and NUR, generating a 3.30 Å map. The resulting 1,217,462 particles were subjected to two rounds of polishing in Relion, HRC, and NUR. The final 908,281 particles were refined to 2.87 Å. Then, the particles were expanded using C3 symmetry and applied to local 3D classification with a mask covering the protomer in Relion. No other conformations were found following symmetry expansion and local 3D classification. For the hEAAT3-X with 10 mM R2-HG dataset, the movie alignments, and micrograph CTF estimation were performed in Relion 4. 3,622,598 particles were auto-picked using template picking and extracted with a box size of 160 pixels (2-fold binning). The particles were imported into CryoSPARC v4 for 2D classification, 2D cleaning, and HRC as the L-Asp dataset. 1,233,807 particles, refined to 3.81 Å, were re-imported into Relion 4 and extracted with a box size of 320 pixels without binning. These particles were further processed as the L-Asp dataset; the final 3.07 Å map was reconstituted using 773,970 particles. Symmetry expansion and local 3D classification performed in CryoSPARC sorted out about 8% of monomers in a minor conformation. For the hEAAT3-X with 10 mM D-Asp dataset, the movies were aligned by MotionCorr2 implemented in Relion 3, and the micrograph CTF parameters were estimated using CtfFfind-4.1. 3,346,010 particles were selected by LoG, extracted with a box size of 256 pixels, and imported into CryoSPARC v3 for 2D classification. 719,954 particles showing secondary features were selected and subjected to *ab initio* reconstruction followed by NUR with C3 symmetry to generate a good template. 3,075,243 particles after 2D cleaning were subjected to two rounds of HRC using one good model and seven decoy volumes. 444,289 particles were selected and refined to 3.29 Å by NUR. After two rounds of polishing in Relion, HRC, and NUR, 391,308 particles were refined to 2.73 Å by NUR with C3 symmetry. Symmetry expansion and local 3D classification did not identify multiple conformations in this dataset. For the hEAAT3-X with L-Cys subset A, 5,756 movies were aligned using MotionCorr2 implemented in Relion 3 with 2-fold binning. The micrograph CTF parameters were estimated using CtfFfind-4.1. 2,538,702 particles were selected using LoG and extracted with a box size of 300 pixels. Particles were imported into CryoSPARC v3 for 2D classification. The good template was generated using particles showing 2D features as previously described. Separately, all the particles after 2D classification were used in *ab initio* reconstruction with less than 10 iterations to generate 7 noise volumes. 2,268,928 particles after 2D cleaning were further cleaned by HRC with one good template and 7 decoy noise volumes. After that, 902,201 particles were reconstituted to 2.8 Å with C3 symmetry by NUR. Then, the particles were re-imported to Relion using PyEM and subjected to Bayesian polishing. The polished particles underwent one round of HRC and NUR to improve resolution. The second round of polishing, HRC, and NUR procedures finally generated a 2.43 Å map with 653,778 particles. Subset B was processed in parallel using a similar strategy. 1,641,561 particles were extracted from 3,757 micrographs and imported into CryoSPARC for 2D classification. After 2D cleaning, 1,474,916 particles underwent further cleaning through heterogeneous refinement. The resulting 614,371 particles were refined to 2.92 Å by NUR with C3 symmetry. After two rounds of polishing in Relion, HRC, and NUR, 444,946 particles were refined to 2.54 Å. 1,112,764 particles from two subsets were combined and refined to 2.36 Å by NUR. These particles were applied to symmetry expansion and local 3D classification. Individual classes of interest were further subjected to local refinement in CryoSPARC.

### Model building and refinement

hEAAT3-X structures with bound L-Glu in iOFS*, hEAAT3-X bound to Na^+^ ions in OFS, and iOFS, and hEAAT3g with bound L-Asp (PDB accession codes: 8CTC, 8CV2 and 8CV3, and 6X2Z respectively) were fitted into EM density maps using ChimeraX^78^. The models were manually adjusted in COOT^79^ and subjected to real-space refinement in Phenix^80^. Structural model validation was performed in Phenix. All the structural figures were prepared using ChimeraX.

## Supporting information

This file contains supplementary figures 1-9 and supplementary table 1.

This movie shows the partial and complete HP2 gate closure upon substrate and Na2 binding

## Acknowledgments

We thank Dr. Xiaoyu Wang, Dr. Qianyi Wu, Dr. Krishna Reddy, and Dr. Yun Huang for the useful discussions. We thank Jing Wang at SEMC-NYSBC, Bing Wang and William Rice at NYU Langone’s Cryo-EM laboratory, and Edwin Fluck at Weill Cornell Cryo-EM facility center for assistance with data collection.

## Author contribution and interest conflict

B.Q. performed the experiments; B.Q. and O.B. conceived the projects, analyzed data, and wrote the manuscript. Competing interests: The authors declare no competing commercial interests.

## Data availability

The Cryo-EM maps and atomic coordinates have been deposited in the Electron Microscopy Data Bank (EMDB) and Protein Data Bank (PDB) under accession code: EMD-46586, PDB-9D66 (hEAAT3-X with L-Asp bound at iOFS*); EMD-46587, (hEAAT3-X in sodium and R-2HG at OFS); EMD-46588, PDB-9D67 (hEAAT3-X with D-Asp bound at iOFS*); EMD-46589, PDB-9D68 (hEAAT3-X with L-Cys bound at OFS, semi-open HP2); EMD-46590, PDB-9D69 (hEAAT3-X with L-Cys bound at iOFS); EMD-46591, PDB-9D6A (hEAAT3-X with L-Cys bound at iOFS*); EMD-46592 (hEAAT3-X in sodium and L-Cys at IFS).

