## Supplementary material for "Structural basis of the excitatory amino acid transporter 3 substrate recognition": This file contains supplementary figures 1-9 and supplementary table 1.

**This PDF file includes the following:**

Supplementary Figures 1-9

Supplementary Movie 1

Supplementary Table1 1

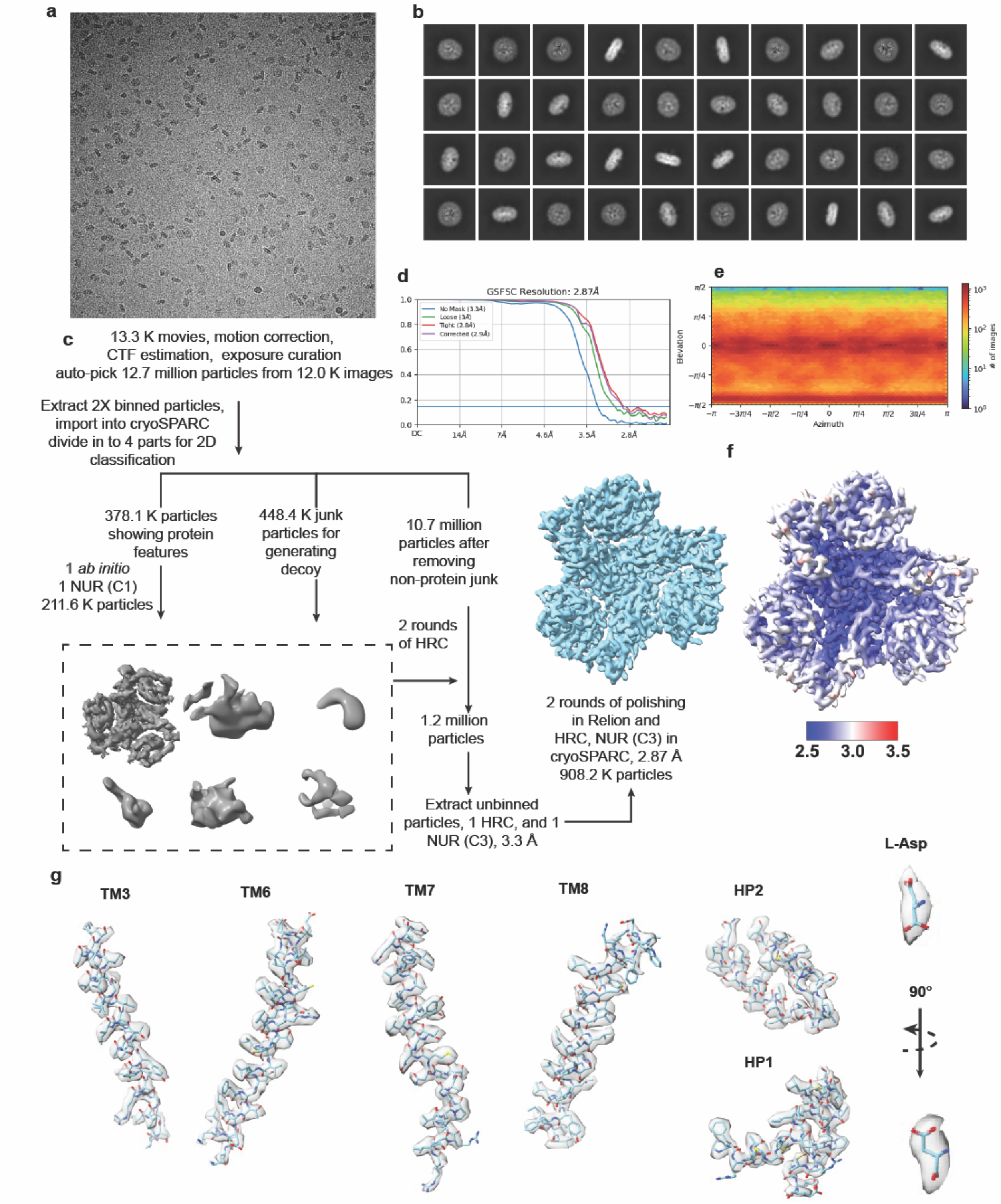


**Supplementary Figure 1: Cryo-EM analysis of hEAAT3-X in complex with L-Asp.** A representative image (**a**) and selected 2D class averages (**b**) of the L-Asp dataset. (**c**), Cryo-EM data processing flow. (**d**), The golden standard Fourier shell correlation (FSC) curves of the final refinement. (**e**), The angular distribution of particles used for the final 3D reconstitutions. (**f**), The local resolution distribution of the final map. (**g**) The EM density of L-Asp, transport domain transmembrane helices (TMs), and helical hairpins (HPs); the map contour level is 0.614 in ChimeraX.

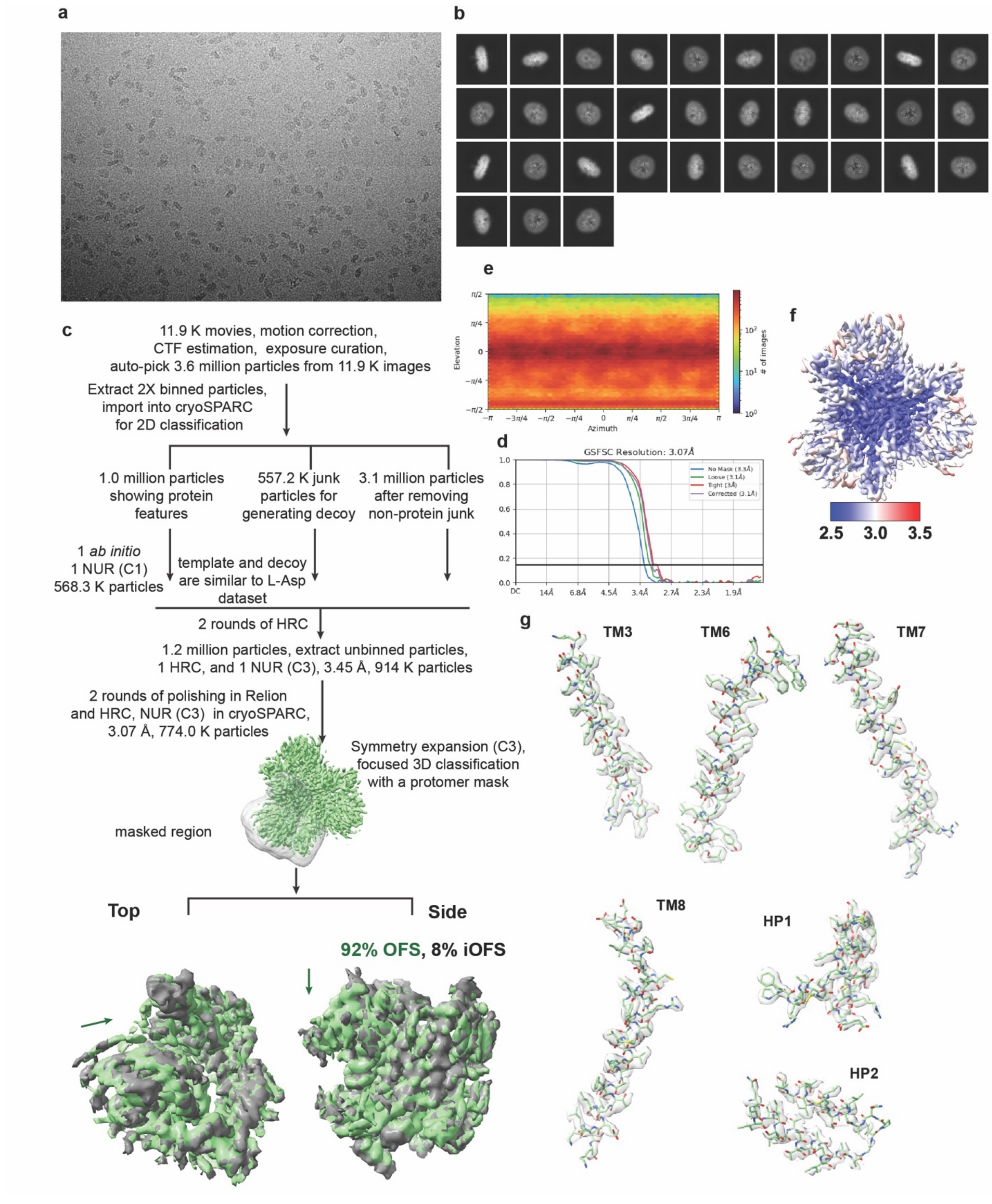
a
b
e
f
2.5
3.0
3.5
c
11.9 K movies, motion correction,
CTF estimation, exposure curation,
auto-pick 3.6 million particles from 11.9 K images
Extract 2X binned particles,
import into cryoSPARC
for 2D classification
1.0 million particles
showing protein
features
557.2 K junk
particles for
generating decoy
3.1 million particles
after removing
non-protein junk
1 ab initio
1 NUR (C1)
568.3 K particles
template and decoy
are similar to L-Asp
dataset
2 rounds of HRC
1.2 million particles, extract unbinned particles,
1 HRC, and 1 NUR (C3), 3.45 Å, 914 K particles
2 rounds of polishing in Relion
and HRC, NUR (C3) in cryoSPARC,
3.07 Å, 774.0 K particles
Symmetry expansion (C3),
focused 3D classification
with a protomer mask
masked region
Top
Side
92% OFS, 8% iOFS
Elevation
Azimuth
### of images
d
GSFSC Resolution: 3.07Å
No Mask (3.3Å)
Loose (3.1Å)
Tight (3Å)
Corrected (3.1Å)
g
TM3
TM6
TM7
TM8
HP1
HP2

**Supplementary Figure 2: Cryo-EM analysis of hEAAT3-X with R-2HG.** A representative image (**a**) and selected 2D class averages (**b**) of the R-2HG dataset. (**c**), Cryo-EM data processing flow; the template and decoy volumes, similar to those in Supplementary Figure 1, are not shown for clarity; the green arrows at the bottom show the transport domain movement from OFS to iOFS. (**d**), The FSC curves of the final refinement. (**e**), The angular distribution of the particles used for the final 3D reconstitutions. (**f**), The local resolution distribution of the final map. (**g**), The EM density of the transport domain TMs and HPs; the map contour level is 0.34 in ChimeraX. The molecular model of $Na^+$-only bound EAAT3-X (PDB: 8CV2) was fitted into the final map without refinement for reference.

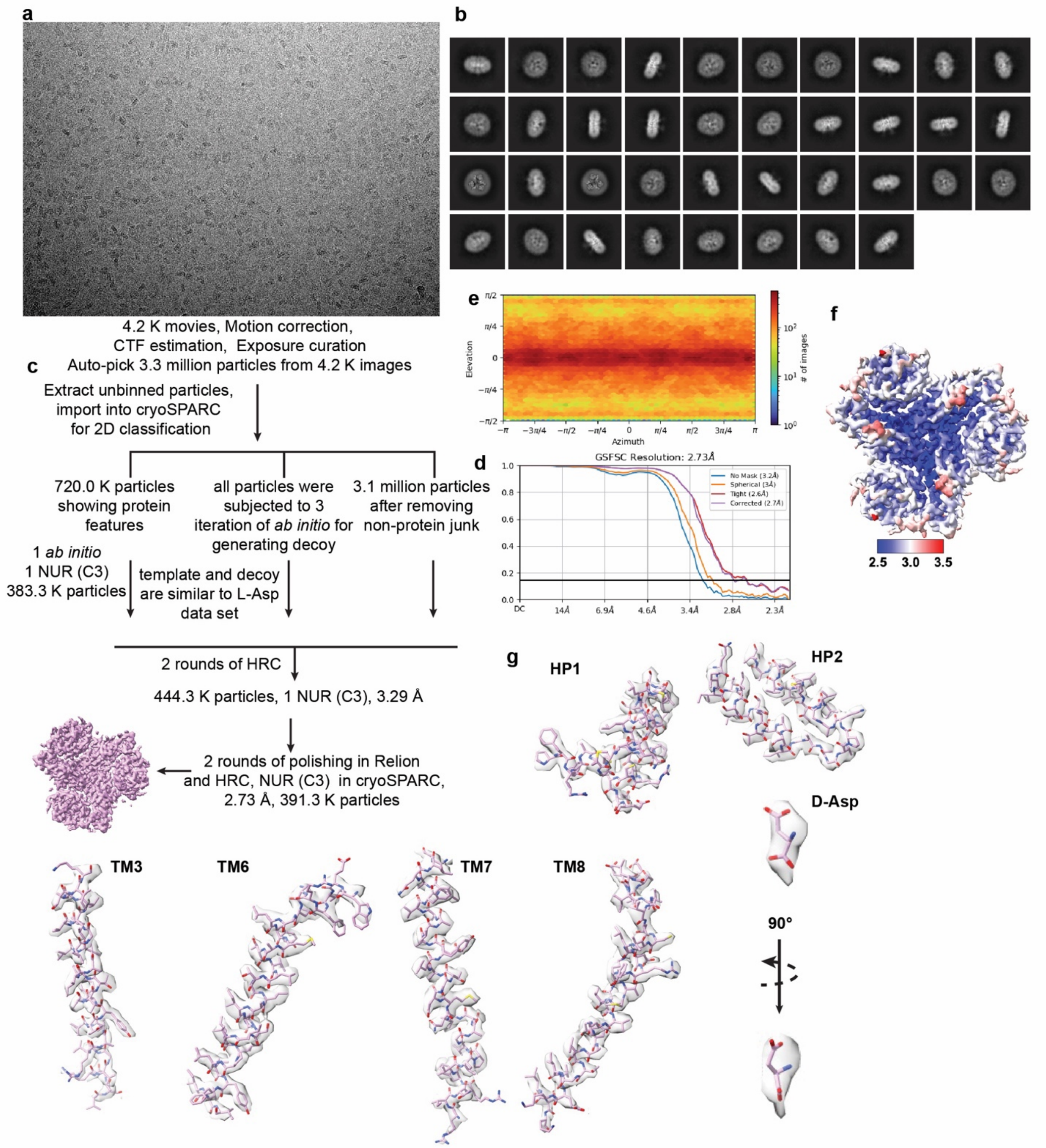


**Supplementary Figure 3: Cryo-EM analysis of hEAAT3-X in complex with D-Asp.** A representative image (**a**) and selected 2D class averages (**b**) of the D-Asp dataset. (**c**), The cryo-EM data processing flow. (**d**), The FSC curves of the final refinement. (**e**), The angular distribution of particles used for the final 3D reconstitutions. (**f**), The local resolution distribution of the final map. (**g**) The EM density of D-Asp, the TMs, and HPs; the contour level is 0.614 in ChimeraX.

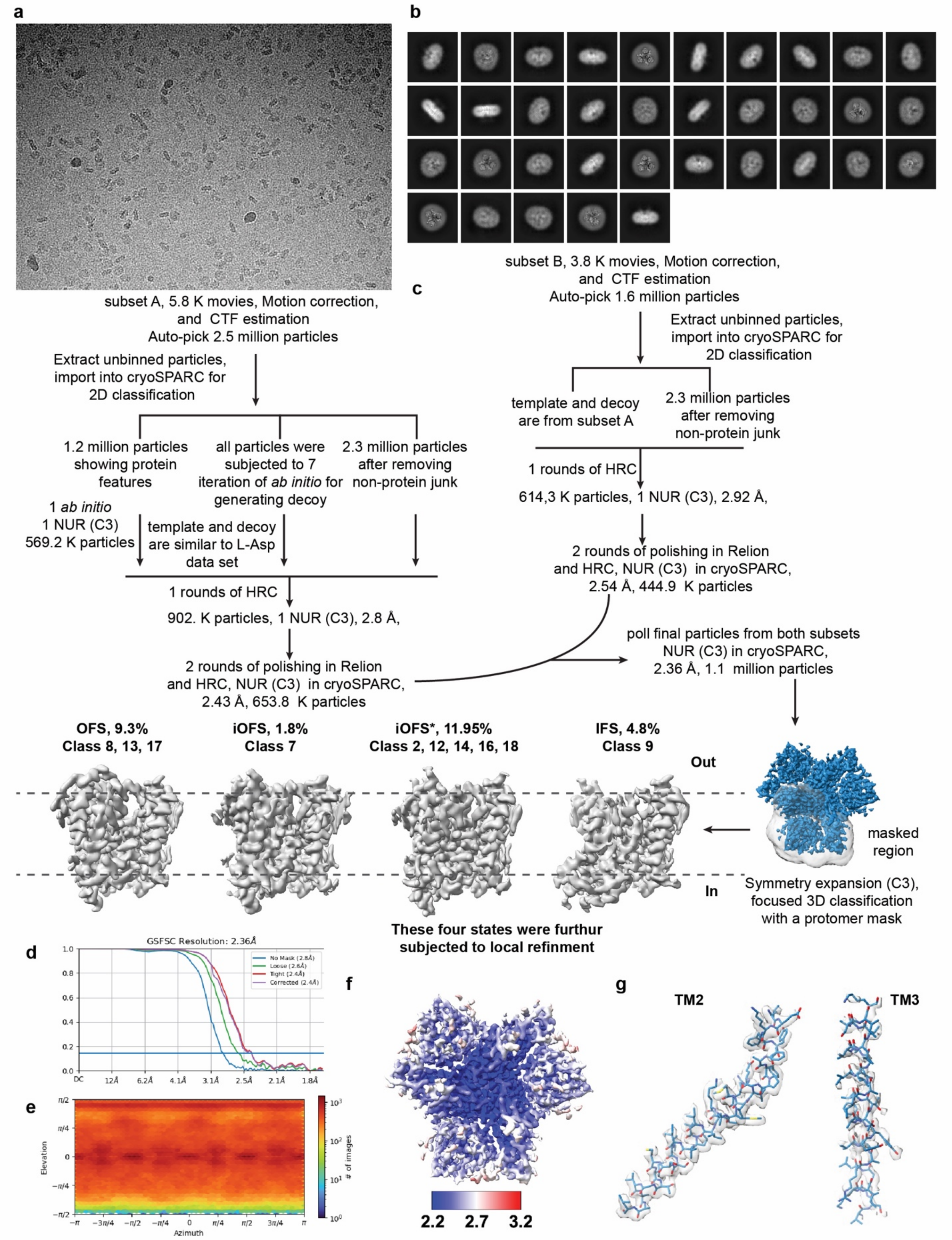
a
b
c
subset A, 5.8 K movies, Motion correction, and CTF estimation
Auto-pick 2.5 million particles
Extract unbinned particles, import into cryoSPARC for 2D classification
1.2 million particles showing protein features
all particles were subjected to 7 iteration of *ab initio* for generating decoy
2.3 million particles after removing non-protein junk
1 *ab initio*
1 NUR (C3)
569.2 K particles
template and decoy are similar to L-Asp data set
1 rounds of HRC
902. K particles, 1 NUR (C3), 2.8 Å,
2 rounds of polishing in Relion and HRC, NUR (C3) in cryoSPARC, 2.43 Å, 653.8 K particles
subset B, 3.8 K movies, Motion correction, and CTF estimation
Auto-pick 1.6 million particles
Extract unbinned particles, import into cryoSPARC for 2D classification
template and decoy are from subset A
2.3 million particles after removing non-protein junk
1 rounds of HRC
614,3 K particles, 1 NUR (C3), 2.92 Å,
2 rounds of polishing in Relion and HRC, NUR (C3) in cryoSPARC, 2.54 Å, 444.9 K particles
poll final particles from both subsets NUR (C3) in cryoSPARC, 2.36 Å, 1.1 million particles
OFS, 9.3%
Class 8, 13, 17
iOFS, 1.8%
Class 7
iOFS*, 11.95%
Class 2, 12, 14, 16, 18
IFS, 4.8%
Class 9
Out
In
masked region
Symmetry expansion (C3), focused 3D classification with a protomer mask
These four states were furthur subjected to local refinment
d
GSFSC Resolution: 2.36Å
No Mask (2.8Å)
Loose (2.6Å)
Tight (2.4Å)
Corrected (2.4Å)
e
Elevation
Azimuth
### of images
f
2.2
2.7
3.2
g
TM2
TM3

**Supplementary Figure 4: Cryo-EM analysis of hEAAT3-X in complex with L-Cys.** A representative image (**a**) and selected 2D class averages (**b**) of the L-Cys dataset. (**c**), Cryo-EM data processing flow. The protomer's reconstitution information is shown in Supplementary Figures 5 and 6. (**d**), The FSC curves of the trimer map. (**e**), The angular distributions of particles used for the 3D reconstitutions of the trimer. (**f**), Local resolution distribution of the trimer map. (**g**) The EM density of TM2 in the scaffold domain and TM3 in the transport domain with the contour level of 0.62 in ChimeraX; the blurred density of TM3 reflects the dynamics of the transport domain.

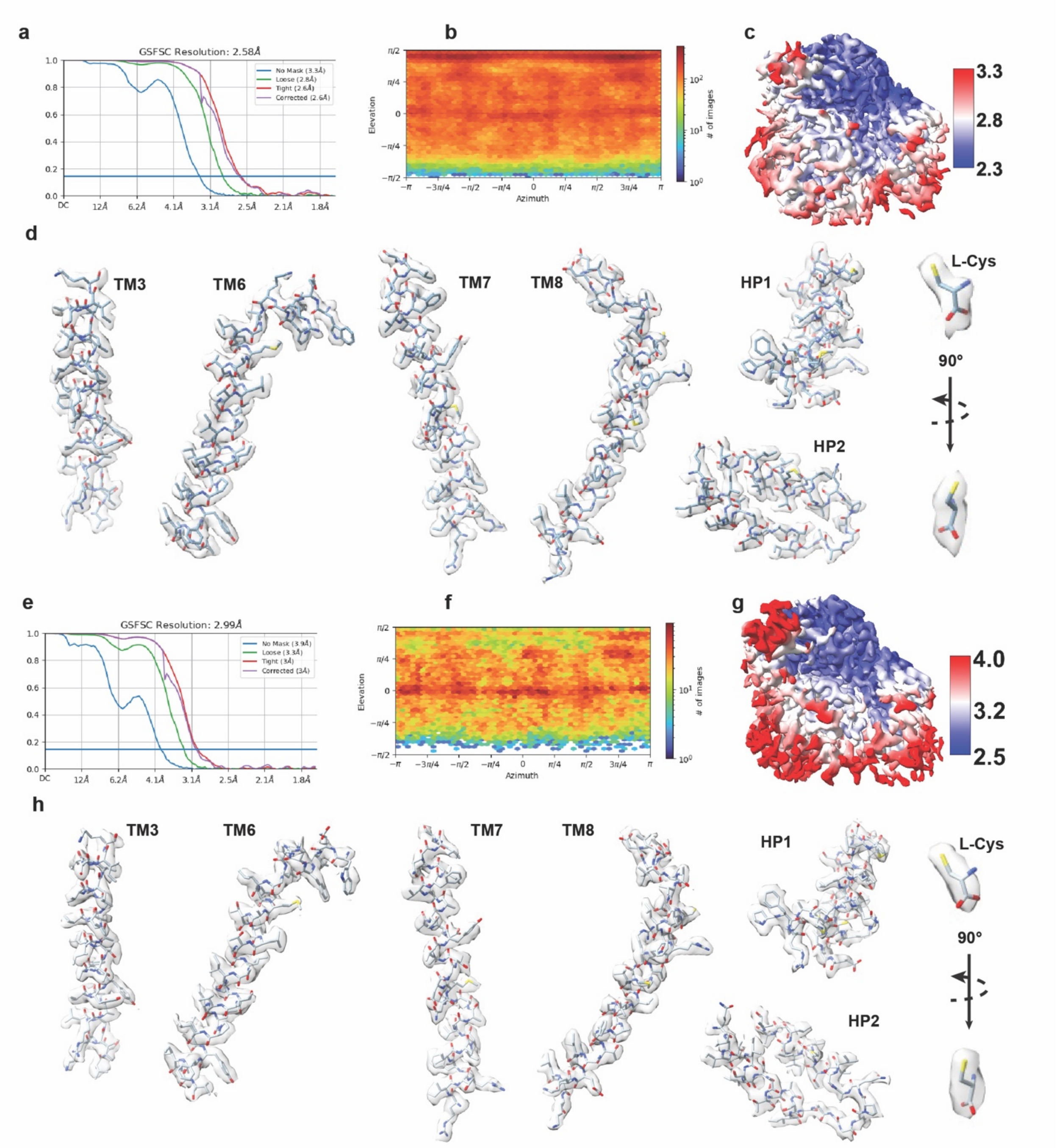


**Supplementary Figure 5: The local refinement of hEAAT3-X in complex with L-Cys in OFS and iOFS.** The FSC curves of the OFS-L-Cys (**a**), and iOFS-L-Cys (**e**). The angular distribution of particles used for the 3D reconstitutions of OFS-L-Cys (**b**) and iOFS-L-Cys (**f**). The local resolution distribution of OFS-L-Cys map (**c**) and iOFS-L-Cys map (**g**). The EM density of L-Cys, transport domain TMs, and HPs of OFS-L-Cys (**d**) and iOFS-L-Cys (**h**). The map contour levels in (**d**) and (**h**) are 0.65 and 0.54, respectively.

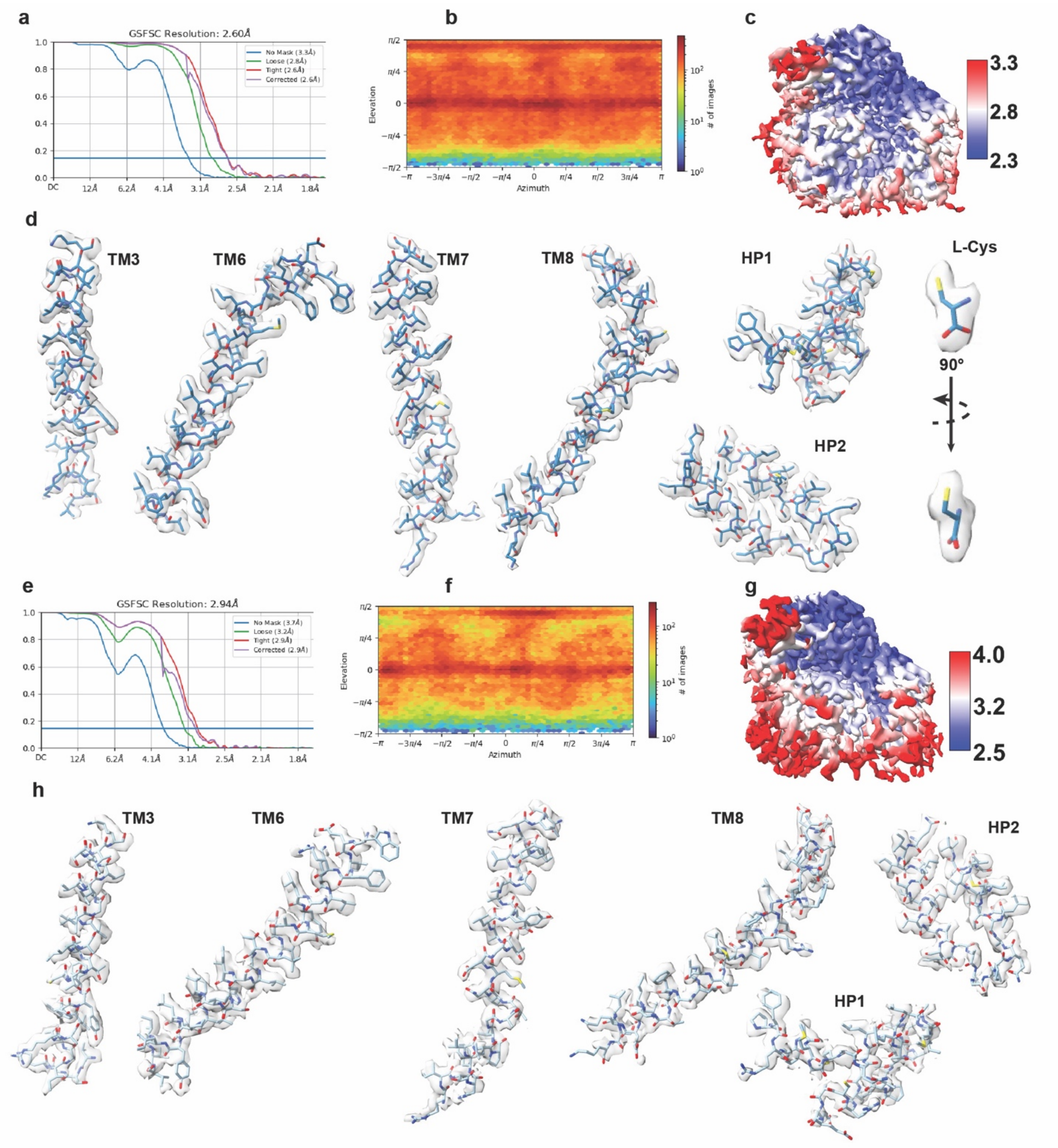


**Supplementary Figure 6: The local refinement of hEAAT3-X in complex with L-Cys in iOFS* and sodium-only bound IFS (IFS-Na$^+$).** The FSC curves of the iOFS*-L-Cys (**a**) and IFS-Na$^+$ (**e**). The angular distribution of particles used for the 3D reconstitutions of iOFS*-L-Cys (**b**) and IFS-Na$^+$ (**f**). The local resolution distribution of iOFS*-L-Cys (**c**) and IFS-Na$^+$ (**g**). The EM density of L-Cys and transport domain TMs and HPs of iOFS*-L-Cys (**d**) and IFS-Na$^+$ (**h**). The map contour levels in (**d**) and (**h**) are 0.61 and 0.43, respectively. The IFS-$Na^+$ molecular model (PDB: 6X2L) was fitted in the IFS-$Na^+$ map without refinement for reference.

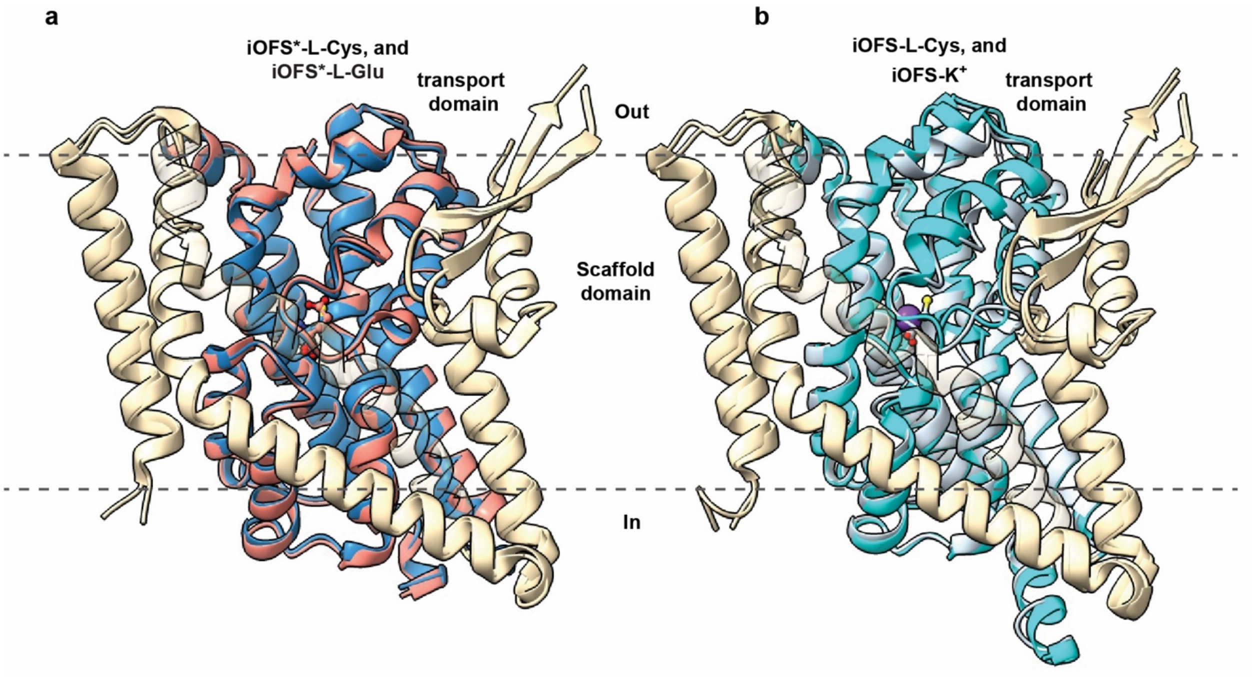


**Supplementary Figure 7: The overall structures of iOFS*-L-Cys and iOFS-L-Cys.** (**a, b**), Superpositions over the entire protomers of (**a**) iOFS*-L-Cys (Blue) and iOFS*-Glu (salmon, PDB: 8CTC) and (**b**) iOFS-L-Cys (light pastel blue) and iOFS-$K^+$ (cyan, PDB: 8CUA). The scaffold domains are colored in wheat, and TM2 is rendered transparent for clarity. The bound L-Glu, L-Cys, and potassium ion are shown as ball-and-stick models and a sphere.

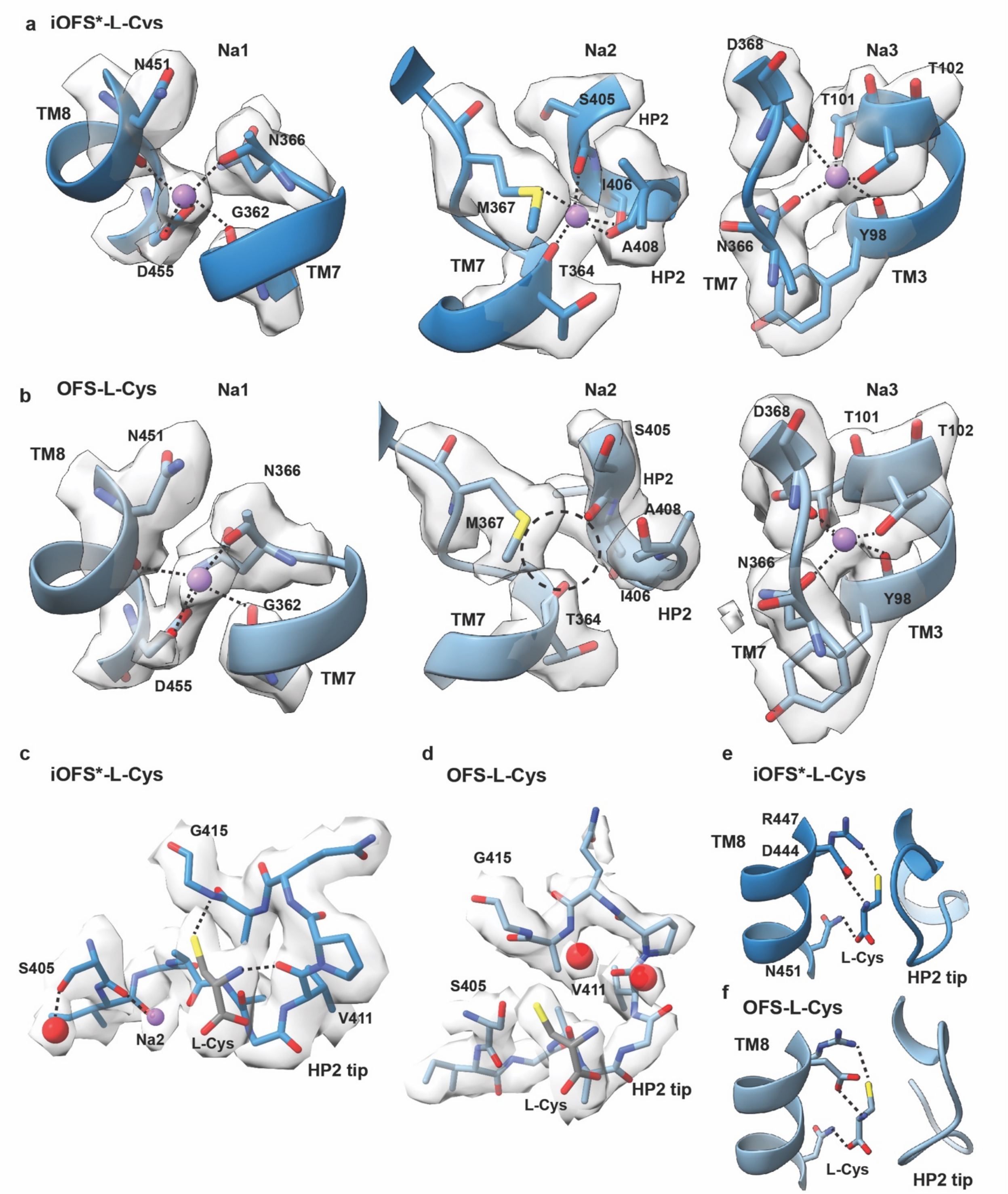


**Supplementary Figure 8: The EM density at sodium-binding sites and the structure of HP2 tip in iOFS*-L-Cys (dark blue) and OFS-L-Cys (pastel blue).** The EM density and geometry of sodium sites in iOFS*-L-Cys (**a**) and OFS-L-Cys (**b**). From left to right: Na1, Na2, and Na3 sites. The dashed circle highlights the distorted Na2 site in OFS-L-Cys. (**c**), The interactions between L-Cys thiolate and the HP2 tip in iOFS*-L-Cys are highlighted as dashed lines. (**d**), They are disrupted in OFS-L-Cys, with water molecules (red spheres) entering the enlarged space. Interactions between L-Cys and coordinating amino acids in TM8 are similar in iOFS*-L-Cys (**e**) and OFS-L-Cys (**f**). $Na^+$ ions and water molecules are shown as spheres scaled to 0.5-fold of their van der Waals radii. The contour levels of density maps in iOFS*-L-Cys and OFS-L-Cys are 0.6 and 0.5, respectively. The dashed black lines show the interaction between protein residues and ligands.

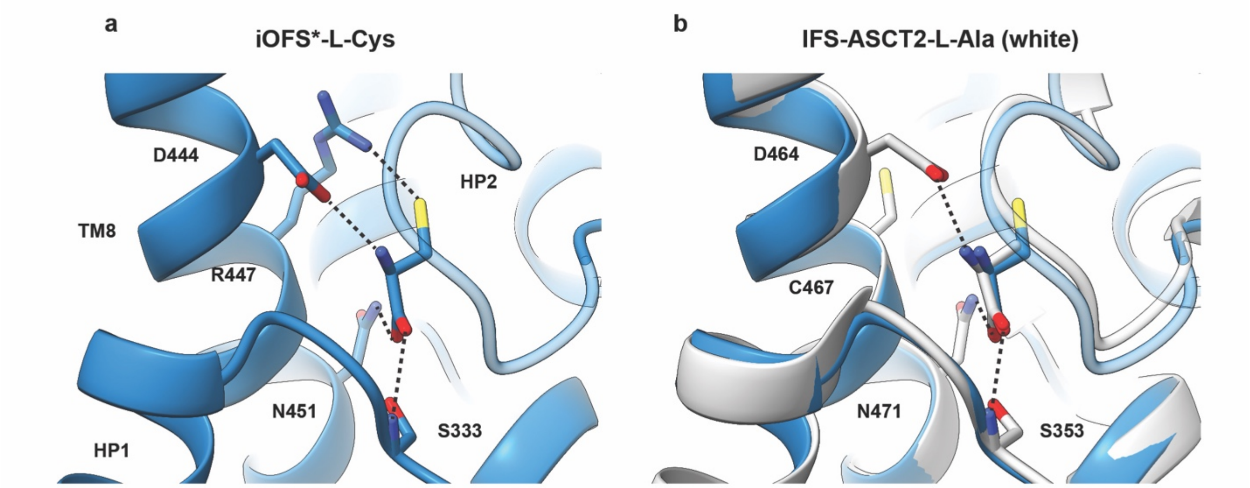


**Supplementary Figure 9: L-Cys recognition in EAAT3 and ASCT2.** (**a, b**), The dashed black lines show the interaction of the residues in TM8 with substrates in EAAT3 (**a,** slate blue), and ASCT2 (**b,** PDB: 8OUD, white). The superposition (**b**) shows nearly identical protein structures and similar poses of bound L-Cys in EAAT3 and L-Ala in ASCT2. C467 in ASCT2 is at the equivalent position to R447 in EAAT3; it is likely too far to interact with bound L-Cys. The structures are superposed on the cytoplasmic halves of their transport domains (residues 314-372 and 442-465 in EAAT and residues 334-392, 462-485 in ASCT2).

**Supplementary Movie 1: The partial and complete HP2 gate closure upon substrate and Na2 binding.** The protein model colors, and coordinate files are the same as in Figure 5. Na1 and Na3 are not shown for clarity. The cytoplasmic halves of the transport domains (residues 314-372 and 442-465) were used for superposition and morphing.

**Supplementary Table 1: Cryo-EM data collection, processing, and model refinement Statistics**

| | iOFS*-L-Asp | OFS-R2HG ($Na^+$ only) | iOFS*-D-Asp | OFS-L-Cys | iOFS-L-Cys | iOFS*-L-Cys | IFS-Cys ($Na^+$ only) |
|---|---|---|---|---|---|---|---|
| **Data collection and processing** | | | | | | | |
| Magnification | 100,100 X | 100,500 X | 64,000 X | 100,500 X | | | |
| Voltage (kV) | 200 | 300 | 300 | 300 | | | |
| Electron exposure (e–/Å$^2$) | 40 | 50.54 | 52.19 | 58.25 | | | |
| Defocus range (μm) | -1.0 ~ -2.5 | -0.8 ~ -2.2 | -0.5 ~ -2.0 | -0.8 ~ -2.4 | | | |
| Pixel size (Å) | 1.16 | 0.844 | 1.076 | 0.825 | | | |
| Symmetry imposed | C3 | C3 | C3 | C3 for initial trimer, C1 for expanded protomer | | | |
| Initial particle images (no.) | 12,668,720 | 3,6262,598 | 3,346,010 | 4,180,263 initial; 1,112,764 final; 3,338,292 C3 expanded | | | |
| Final particle images (no.) | 908,281 | 773,970 | 391,308 | 307,042 | 60,670 | 365,660 | 159,538 |
| Map resolution (Å) | 2.87 | 3.07 | 2.73 | 2.58 | 2.99 | 2.60 | 2.94 |
| FSC threshold | 0.143 | 0.143 | 0.143 | 0.143 | 0.143 | 0.143 | 0.143 |
| Map resolution range (Å) | 7.35 ~ 2.56 | 7.19 ~ 2.58 | 30.17 ~ 2.38 | 41.24 ~ 2.31 | 38.18 ~ 2.65 | 30.80 ~ 2.35 | 42.46 ~ 2.58 |
| **Refinement** | | This map is similar to EMD-27006 (EAAT3-X, OFS-$Na^+$)., The model 8CV2 can be fitted in this map. | | | | | This map is similar to EMD-22011 (EAAT3, IFS-$Na^+$). The model 6X2L can be fitted in this map. |
| Initial model used (PDB code) | 8CTC | | 8CTC | 6X2Z | 8CV3 | 8CTC | |
| Model resolution (Å) | 3.1 | | 3.0 | 2.8 | 3.3 | 2.7 | |
| FSC threshold | 0.5 | | 0.5 | 0.5 | 0.5 | 0.5 | |
| Map sharpening *B* factor (Å$^2$) | -163 | | -107 | | | | |
| Model composition | | | | | | | |
| Non-hydrogen atoms | 9,306 | | 9,297 | 3,242 | 3,143 | 3,105 | |
| Protein residues | 1,221 | | 1,218 | 426 | 413 | 407 | |
| Ligands | 15 | | 15 | 3 | 5 | 5 | |
| *B* factors (Å$^2$) | | | | | | | |
| Protein | 51.23 | | 65.75 | 40.46 | 59.17 | 38.93 | |
| Ligand | 88.27 | | 82.08 | 41.15 | 104.00 | 70.93 | |
| R.m.s. deviations | | | | | | | |
| Bond lengths (Å) | 0.005 | | 0.004 | 0.004 | 0.004 | 0.004 | |
| Bond angles (°) | 1.078 | | 0.979 | 0.934 | 0.953 | 0.904 | |
| Validation | | | | | | | |
| MolProbity score | 1.13 | | 1.27 | 1.12 | 1.17 | 1.10 | |
| Clashscore | 3.35 | | 4.81 | 3.30 | 3.87 | 3.14 | |
| Poor rotamers (%) | 0.00 | | 0.00 | 0.00 | 0.00 | 0.00 | |
| Ramachandran plot | | | | | | | |
| Favored (%) | 98.25 | | 97.92 | 99.05 | 98.28 | 98.25 | |
| Allowed (%) | 1.75 | | 2.08 | 0.95 | 1.72 | 1.75 | |
| Disallowed (%) | 0.00 | | 0.00 | 0.00 | 0.00 | 0.00 | |
| **PDB code** | 9D66 | | 9D67 | 9D68 | 9D69 | 9D6A | |
| **EMDB code** | 46586 | 46587 | 46588 | 46589 | 46590 | 46591 | 46592 |
